# Driving theta-gamma oscillations modulates extrasynaptic GABAergic tone: a tACS-TMS study

**DOI:** 10.64898/2026.01.26.700604

**Authors:** Camille Lasbareilles, Valentina Mancini, Alek Pogosyan, Helen Zhang, Charlotte Austin, Huiling Tan, Charlotte J Stagg

## Abstract

**BACKGROUND:** Theta-gamma phase-amplitude coupled (θγ-PAC) oscillations in primary motor cortex (M1) have been shown to support motor skill acquisition. Past research has shown that driving gamma activity at the theta peak (TGP), but not the theta trough (TGT) using transcranial alternating current stimulation (tACS) enhances motor learning (Akkad et al., 2021). However, the neurophysiological mechanisms underlying this phase-specific effect remain unclear.

**METHODS:** In a double-blind, sham-controlled, cross-over study, twenty-two healthy participants received 20 minutes of 75Hz/6Hz TGP-tACS, TGT-tACS, or sham stimulation over M1. We used paired-pulse transcranial magnetic stimulation (TMS) to assess GABAergic and NMDAR-mediated activity before, during, and after tACS. Outcome measures included short-interval intracortical inhibition at 1ms (SICI1ms; extrasynaptic GABAergic tone) and 2.5ms (SICI2.5ms; synaptic GABAA activity), intracortical facilitation at 12ms (ICF12ms; NMDAR activity), and motor evoked potential (MEP) amplitude (corticospinal excitability).

**RESULTS:** TGP-tACS selectively decreased SICI1ms, a putative marker of extrasynaptic GABAergic tone (main effect of Stimulation: p=.021), with significant differences between TGP and TGT during late stimulation (p=.047). No significant effects were observed on corticospinal excitability, synaptic GABAergic activity (SICI2.5ms), or NMDAR signalling (ICF12ms).

**CONCLUSIONS:** Driving theta-gamma oscillations at the theta peak using tACS specifically modulates extrasynaptic GABAergic tone in M1 without affecting corticospinal excitability or synaptic inhibition. Given that reductions in GABAergic signalling supports motor learning, these findings provide a neurophysiological mechanism for the phase-specific behavioural effects of θγ-PAC tACS and suggest a potential therapeutic approach for facilitating motor recovery after stroke.

**Highlights:**

- Theta-gamma peak tACS selectively reduces extrasynaptic GABA in human motor cortex
- Only gamma at the peak, not trough, of theta stimulation modulates GABA
- No effects on corticospinal excitability, synaptic GABA, or NMDAR signalling
- tACS-TMS reveals mechanism for phase-dependent motor learning effects

## Introduction

In rodent hippocampus, gamma frequency activity (γ; 30-100Hz) coupled to theta (θ; 4-6Hz), is critical for learning^1,2^. In humans, cross-frequency, phase amplitude coupled θγ oscillations (θγ-PAC) occurs in multiple brain regions to support numerous computations. Notably, in primary motor cortex (M1), γ-activity observed at the theta peak^3^ is related to motor skill aquisition^4,5^.

Artificially entraining θγ-PAC using transcranial alternating current stimulation (tACS) in M1 improves motor behaviour, but only when γ-activity is coupled to the positive half [θγ peak; TGP], not the negative half [θγ trough; TGT] of a θ-oscillation^4^. Given that motor learning depends on GABAergic signalling changes^6^, TGP-tACS may modulate GABAergic signalling to support skill acquisition. However, the underlying neurophysiological mechanisms that govern the phase-specificity of θγ dynamics on motor behaviour remain unknown.

We tested the hypothesis that TGP-tACS modulates GABAergic activity. Using a combined tACS-TMS approach, we assessed the on- and off-line effects of driving θγ-PAC (TGP and TGT) oscillations on proxies for GABA- and NMDA_R_-mediated changes in excitation/inhibition using paired-pulse TMS.

## Methods

Twenty-two healthy, right-handed participants (12 female, aged 18-35) provided written informed consent for this double-blind, sham-controlled, cross-over study (ethical approval R81071/RE002). tACS was applied via two 5x5cm tES rubber electrodes centred over C3 and Pz^4^.

Participants completed three counter-balanced sessions, at least seven days apart, receiving 20 minutes of 2mA peak-to-peak 75Hz/6Hz TGP, TGT, or sham (two 10 second blocks of 6Hz) tACS (Signal, CED; Figure 1A) via a NeuroConn neurostimulator (Neurocare, Germany). Bang’s Blinding Index for sham was 0.5.

**Figure 1:**
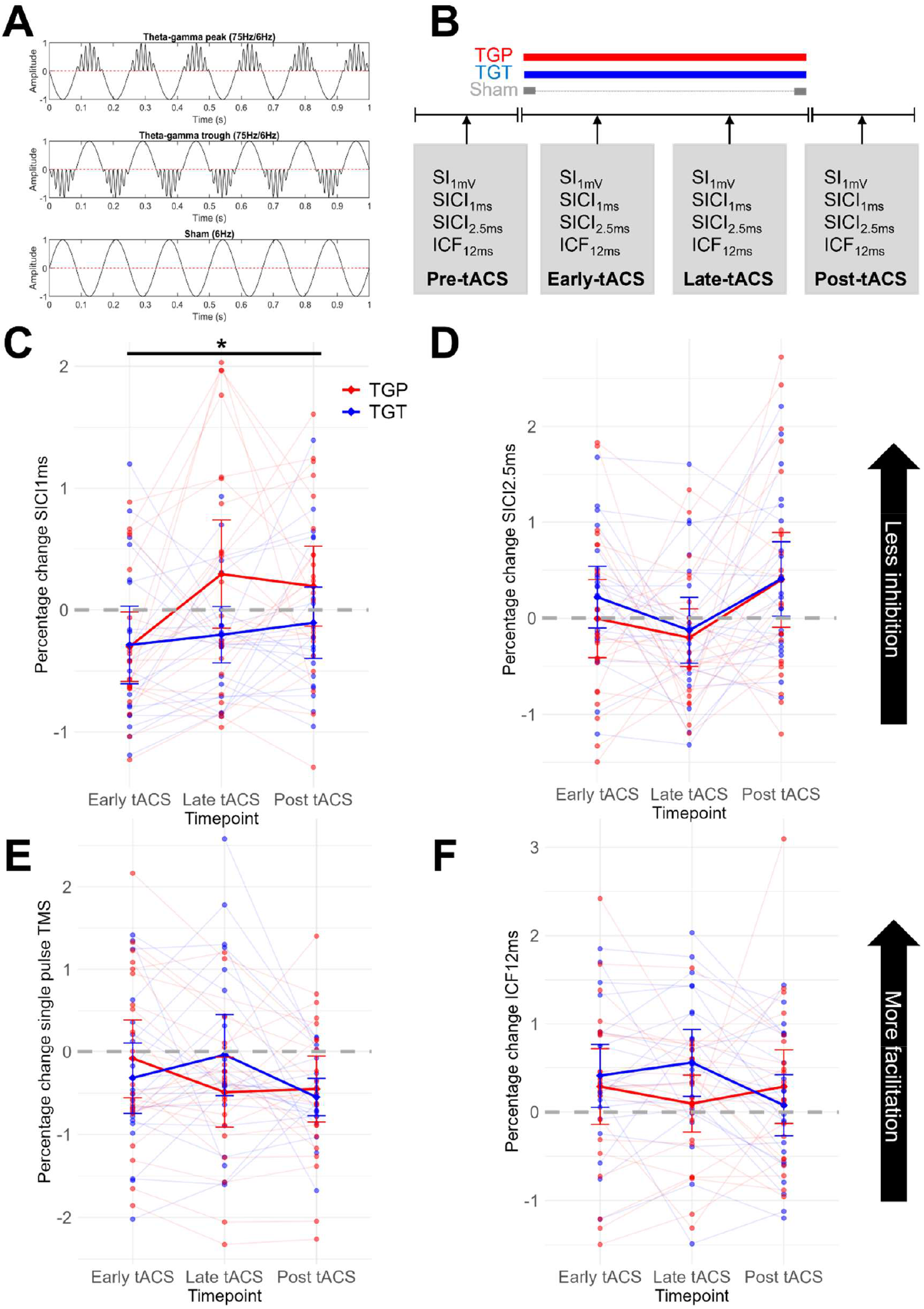
A) One second of the custom-coded TGP, TGT, and sham waveforms delivered. B) Experimental design. SICI: short interval intracortical inhibition. ICF: Intracortical Facilitation. Panels C-F show the percentage change in MEP amplitude from pre tACS (normalised to sham) for C) SICI1ms, D) SICI2.5ms, E), corticospinal excitability (single pulse TMS), and F) ICF12ms. We show that artificially driving motor cortical TGP oscillations using tACS uniquely decreases extrasynaptic GABAergic tone (C) with no significant effect on cortical excitability, synaptic GABAergic tone (SICI2.5ms) or NMDA_R_ signalling (ICF12ms).

Single (spTMS) and paired-pulse TMS (ppTMS) metrics were acquired pre-tACS, during-tACS (*early-* and *late-*tACS), and post-tACS using a Magstim Bistim TMS unit with a standard figure-of-eight 70mm coil (Magstim Company, UK). TMS pulses were applied over the C3 electrode with the TMS coil handle pointing posteriorly and laterally to the mid-sagittal line.

EMG data were recorded from the right first dorsal interosseous muscle (FDI) in a belly-tendon montage, amplified (Digitimer D360; CED) and sampled at 5kHz (1401; CED). We defined SI_1mV_ as the lowest stimulation intensity able to elicit MEPs with a mean peak-to-peak amplitude of 1mV over 10 trials in the relaxed FDI. Active motor threshold (aMT) was defined as the lowest stimulation intensity able to elicit MEPs with a mean peak-to-peak amplitude of ≥0.2mV in 5/10 trials during ∼30% maximum voluntary FDI contraction.

In each TMS block, we acquired *1)* 10 spTMS pulses at 100% SI_1mV_, and *2)* ppTMS measures with interstimulus intervals of 1ms (short interval intracortical inhibition; SICI_1ms_), 2.5ms (SICI_2.5ms_), and 12ms (intracortical facilitation; ICF_12ms_; Figure 1B), with the conditioning stimulus (CS) set to 70% aMT and the test stimulus (TS) to SI_1mV_. The intensity of the TS was adjusted per TMS block to maintain a mean 1mV TS for the ppTMS metrics (SI_1mV_adj_). For each ppTMS metric we calculated the ratio of the mean amplitude of the conditioned and unconditioned MEPs (ppMEP/SI_1mV_adj_).

## Results

We first ensured that there were no differences between sessions at pre-tACS. Repeated-measures ANOVAs on the pre-tACS data with one factor of **Stimulation** (TGP, TGT, sham) showed no significant main effect of **Stimulation** for any TMS protocol (all p >0.039). All ppTMS protocols induced expected inhibition (SICI_1ms_, SICI_2.5ms_) or facilitation (ICF_12ms_) effects at baseline.

Next, we calculated the percentage change in each MEP metric from that session’s mean pre-tACS value (i.e. ([timepoint-pre-tACS]/pre-tACS)x100)). We calculated the absolute change in MEP amplitude from pre-tACS. All data were z-transformed and outliers (+/-3SD from the mean of each stimulation condition at each timepoint) were removed. Active stimulation data were normalised to the mean of the sham at each timepoint. Statistical analyses were performed using the linear mixed effects model (lme4) package in R, with Stimulation (TGP, TGT, sham) and Timepoint (early-tACS, late-tACS, post-tACS) as fixed effects, a Stimulation×Timepoint interaction, and a random intercept of Subject.

θγ-PAC tACS had no significant effect on M1 corticospinal excitability, as quantified by the SI_1mV_ MEP amplitude (no significant main effect of Stimulation (F(1,105.16)=0.146,p=.703) or Timepoint (F(2,105.33)=1.816,p=.168), and no significant Stimulation×Timepoint interaction (F(2,105.56)=1.915,p=.152; Figure 1E)).

Next, we investigated the effect of θγ-PAC tACS on SICI_1ms_, a putative measure of extrasynaptic GABAergic tone. There was a significant main effect of **Stimulation** (F(1,107.40)=5.458,p=.021) and **Timepoint** (F(2,106.83)=3.504,p=.034), but no significant Stimulation×Timepoint interaction (F(2,106.87)=1.3203,p=.271); Figure 1C). Exploratory t-tests revealed a significant difference in percentage change in SICI_1ms_ between TGP and TGT at late-tACS (t(31.26)=2.068,p=.047).

Finally, θγ-PAC tACS had no significant effect on either GABA_A_ synaptic activity, as indexed via SICI_2.5ms_, or NMDA_R_ activity, as indexed via ICF_12ms_ (SICI_2.5ms_: significant main effect of Timepoint [F(2,107.28)=4.062,p=.020], no significant main effect of Stimulation [F(1,107.27)=2.000,p=.160], no significant Stimulation×Timepoint interaction [F(2,107.25)=0.566,p=.569], see figure 1D. ICF_12ms_: no significant main effect of Stimulation [F(1,109.01)=1.118,p=.293], Timepoint [F(2,109.01)=0.772,p=.465] and no significant Stimulation×Timepoint interaction [F(2,109.01)=2.451,p=.091], see Figure 1F).

## Discussion

Overall, artificially driving motor cortical TGP oscillations using tACS uniquely decreased a putative metric of extrasynaptic GABAergic tone (as indexed via SICI_1ms_) with no significant effect on cortical excitability, synaptic GABAergic tone (SICI_2.5ms_), or NMDA_R_ signalling (ICF_12ms_). Given the role of reductions in GABAergic signalling to support motor skill acquisition, this supports previously reported behavioural findings^4^.

We previously showed that driving continuous M1 γ-tACS resulted in a significant, duration-dependent reduction in local synaptic GABA_A_ activity as quantified by SICI_2.5ms_ during movement preparation, with no significant effect on corticospinal excitability^7^. While SICI_2.5ms_ reflects GABA_A_ synaptic activity, SICI_1ms_ has been suggested to reflect extrasynaptic GABAergic tone^8^. Oscillations reflect the coordination of numerous microcircuits, and such oscillations crucially depend on the conduction speed within these microcircuits. Extrasynaptic tone, particularly that determined by GABAergic conductance, plays a key role in determining conduction speed^9^, and hence the circuit oscillations are likely more sensitive to modulations in extrasynaptic tone than phasic inhibition. This link between γ and GABAergic tone is consistent with neuropsychiatric conditions such as schizophrenia, where abnormalities in genes that regulate extrasynaptic GABAergic tone significantly reduce γ-oscillations^10^.

Selective reductions of extrasynaptic GABAergic signalling by TGP tACS support a neurophysiological link between inhibitory activity and γ-activity underpinning learning. Modulating γ-activity therefore represents a potential approach to facilitate motor learning as part of post-stroke recovery.

## Acknowledgements

Conceptualisation: CL, VM, CJS; Investigation: CL, VM, HZ, CA; Methodology: AP, HT; Formal Analysis: CL, HZ, CA; Supervision: HT, CJS; Funding Acquisition: HT, CJS; Writing - original draft: CL; Writing - review and editing: CL, VM, AP, HZ, CA, HT & CJS.

